# Back to the Wild: Polluted Site Remediation and Biosphere Resilience

**DOI:** 10.1101/2021.04.22.441002

**Authors:** Débora Toledo Ramos, Henry Xavier Corseuil, Timothy M. Vogel

## Abstract

Worldwide efforts to depollute environments altered by human industrial activity have begun to produce an ever-increasing number of “clean” sites. “Clean” is defined by local regulatory processes and often responds to low compound concentrations or risk evaluations. Yet, these sites have been critically derailed from their historical biological activity by both the pollution event and the clean-up technology. This work explored the impact of contaminated (and remediated) sites on local microbial ecosystems. Different parcels of the same field site with the same relatively uniform microbial ecology were polluted and cleaned-up over the last 15 years. The statistical evaluation of the perturbation described changes to the local ecosystem that went back to the original baseline microbial composition although the pollution sources and the clean-up technologies affected the rate of return to the pre-disturbed condition. This rate reflected the intensity of the clean-up treatments. The role played by microbial communities on ecosystem maintenance and mitigation of pollution events lays the groundwork for predicting the microbial community responses to perturbations and the ability to reassert themselves. Predictions of ecosystem response to anthropogenic impacts could support decision-making on environmental management strategies for contaminated sites clean-up, depending on the ecosystem services desired to maintain or the risk posed to sensitive receptors.

## 1. INTRODUCTION

### Surviving the wilderness

The ability of microorganisms to resist perturbation and reorganize is termed resilience and reflects the capacity of system self-organization and adaptation to a desirable ecosystem state^1^. Re-establishing the original microbial communities is required to maintain physiological and metabolic processes that are important for ecosystem services (*i.e*. water and soil quality)^2^. Nevertheless, resiliency responses may not be necessarily beneficial to all systems as their perceived benefit depends on the desired ecosystem function^3,4^. For example, if the system is highly resilient and a supplementary compound is added to stimulate nitrifying microorganisms to enhance soil quality, then resilience will not be helpful as it hinders the growth of the new desired community. However, if microbial communities respond to a fuel spill and they start degrading pollutants, the ecosystem will be perturbed. The ecosystem could recover after the perturbation is alleviated, thus contributing to the overall system resilience. Therefore, the dynamics of microbial response to environmental changes must be framed within the continuum of background - contamination - remediation – recovery. Worldwide efforts to depollute environments altered by human industrial activity have begun to produce an ever-increasing number of “clean” sites. “Clean” is defined by local regulatory processes and often responds to low compound concentrations or risk evaluations. Yet, these sites have been critically derailed from their historical biological activity by both the pollution event and, in many cases, the clean-up technology. The long-term effects of our changing climate are also raising questions about critical ecosystem functions^5,6^, which underscores the need to explore the impact of contaminated (and remediated) sites on local ecosystems.

Anticipating the ecosystem response to pollution events perturbations could be the key for supporting environmental management and engineering decisions relative to contaminated sites clean-up^5–7^ These perturbations can affect members of the microbial community in different ways. Ecotoxicological effects might negatively influence some community members, while others might be positively influenced especially if the contaminant provides resources for microbial growth and selects for underrepresented species, which would thrive in the modified environment. In addition to the alterations promoted by the pollutants, the new scenario established by the subsequent pollution treatment using physical, chemical or even biological technologies might also push the ecosystem further from its original structure and function. Unfortunately, to date, no systematic framework of multiple field-based contamination events has been performed due to (i) the difficulties in assessing original baseline microbial composition and post-disturbance composition^5,8^, (ii) the heterogeneity of the background microbial ecology of distant unrelated sites and (iii) the lack of efficient next generation sequencing (NGS) tools to investigate these systems at different genomic resolution.

Our study addressed these knowledge gaps by providing an integrative framework of geochemical processes and microbial composition at undisturbed (background), disturbed (polluted) and post-disturbed (remediated) environments. Microbial response to different pollution sources and clean-up technologies was assessed by identifying pollution bioindicators, resilient communities and the associated time taken to recover to the pre-disturbed condition. The role played by microbial communities on ecosystem maintenance and mitigation of pollution events exemplifies our improved understanding and lays the groundwork for predicting the microbial community response. Predictions of ecosystem response to anthropogenic impacts could help decision-making on environmental management strategies.

## 2. MATERIALS AND METHODS

Different parcels of the same field site with the same relatively uniform microbial ecology were polluted and cleaned-up over the last 15 years (Figure 1) ^9–17^ A range of pollution events was used including pollution by gasoline-ethanol (E10, E25, E85), biodiesel (B100) and diesel-biodiesel blends (B20). Treatment technologies included chemical oxidation (partial chemical oxidation via modified Fenton’s reaction) and biological methods (monitored natural attenuation and biostimulation) and ranged from polluted to relatively successful and completely successful with the pollutant concentrations dropping below detection levels. The different pollution events and clean-up technologies were evaluated relative to background microbial ecosystem communities by high throughput sequencing and by chemical analyses of the different pollutants. The taxonomic profiles of the different samples from the area were determined by sequencing part (V3-V4 regions) of the 16S rRNA gene in the extracted environmental DNA. Chemical analyses (BTEX, PAH, ethanol and methane) were performed by gas chromatography. A detailed description of materials and methods is described below.

**Figure 1.**
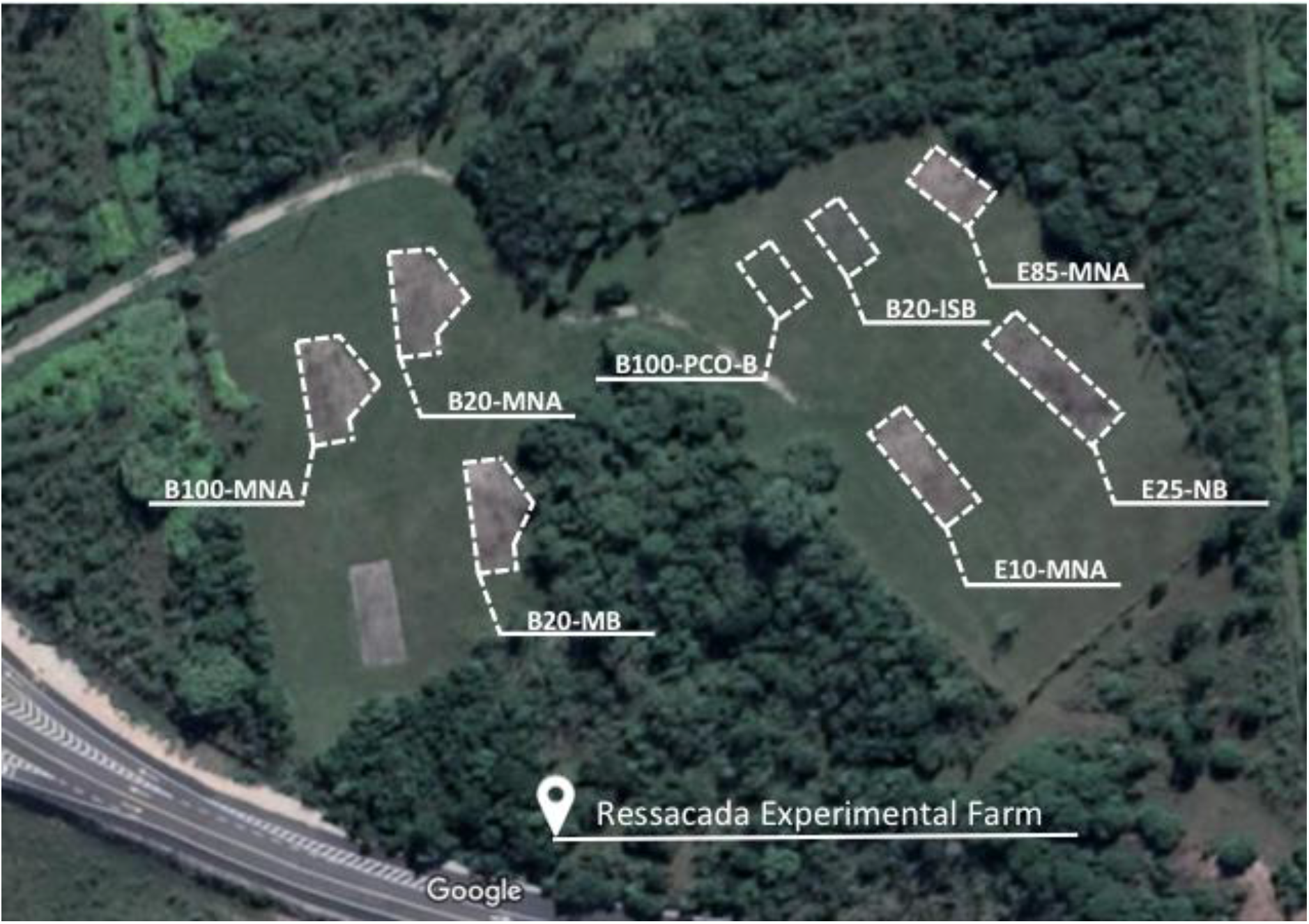
Plan view of the field studies monitored at Ressacada Experimental Farm.

### 2.1 Controlled release field experiments

Field experiments were established by the controlled release of different fuels into source-zone areas of approximately 1 x 1 x 1.5-1.8 m deep down to the water table. This study comprised the following experiments: B100-MNA: pure soybean biodiesel under monitored natural attenuation; B20-MNA: soybean biodiesel and diesel blend (20:80 v/v) under monitored natural attenuation; B20-MB: soybean biodiesel and diesel blend (20:80 v/v) under methanogenic biostimulation; B100-PCO-B: pure palm biodiesel under partial chemical oxidation coupled with biodegradation; B20-ISB: palm biodiesel and diesel blend (20:80 v/v) under combined iron and sulfate biostimulation; E85-MNA and E10-MNA: ethanol and gasoline (85:25 and 10:90, v/v/, respectively) under monitored natural attenuation. A summary of the experiments configuration is provided in Table 1.

**Table 1.** Field experiments configuration, including fuel blends, clean-up technologies and the corresponding monitoring period.

| Experiment | Clean-up Technology | Volume released (L) | Monitoring period |
| --- | --- | --- | --- |
| E25 | NB | 100 | 2004 - 2015 |
| B100 | MNA | 100 | 2008 - 2015 |
| B20 | MNA | 100 | 2008 - 2015 |
| E10 | MNA | 100 | 2009 - 2015 |
| B20 | MB | 100 | 2010 - 2015 |
| E85 | MNA | 200 | 2010 - 2015 |
| B100 | PCO-B | 100 | 2014 - 2015 |
| B20 | ISB | 100 | 2014 - 2015 |

*Note.* MNA. Monitored natural attenuation; MB. Methanogenic biostimulation; PCO-B. partial chemical

### 2.2 Groundwater chemical analysis

Samples were collected from monitoring wells at varying depths (2, 3, 4, 5 and 6 m below ground surface). A peristaltic pump and Teflon tubing were used to collect groundwater samples into capped sterile vials without headspace. BTEX, ethanol and methane were analyzed by gas chromatography using a GC HP model 6890 II equipped with a flame ionization detector (FID), HP 1 capillary column (30 cm, 0.53 mm, 2.65 mm) and HP 7694 headspace auto sampler, as described elsewhere^17^. PAH were extracted from groundwater using solid phase SPE cartridges, according to EPA method 525.2, and measured by gas chromatography (HP model 6890 II with a flame ionization detector (FID) and HP-5 capillary column), as described elsewhere^11^. The detection limits were (in parenthesis): BTEX (1 μg L^−1^ for each), ethanol (1 mg L^−1^) and methane (10 μg L^−1^)^16^. Detection limits were (in parenthesis): naphthalene (7 μg L^−1^), methylnaphthalene (5 μg L^−1^), dimethylnaphthalene (7 μg L^−1^), acenaphthylene (8 μg L^−1^), acenaphthene (8 μg L^−1^), fluorene (8 μg L^−1^), phenanthrene (9 μg L^−1^), anthracene (9 μg L^−1^), fluoranthene (10 μg L^−1^), pyrene (9 μg L^−1^), benzo[a]anthracene (9 μg L^−1^), chrysene (10 μg L^−1^), dibenzo[a,h]anthracene (12 μg L^−1^), benzo[b]fluoranthene (12 μg L^−1^), benzo[k]fluoranthene (31 μg L^−1^), benzo[a] pyrene (36 μg L^−1^), indene [1,2,3-cd]pyrene (28 μg L^−1^) and benzo[g,h,i]pyrene (11 μg L^−1^)^11^.

### 2.3 Microbial analysis

Groundwater samples were filtered with Millipore membrane (polyethersulfone, hydrophilic), 0.22 μm pore size. DNA was extracted according to the MoBio Power Soil kit (Carlsbad, CA) protocol^10^. The 16S ribosomal RNA gene sequencing was performed to assess microbial communities from the eight different field experiments (Figure 1). The variable regions, V3 and V4, of the gene that codes for the 16S rRNA were amplified by PCR and sequencing was performed using Illumina MiSeq technology. The following stepwise procedures were originally described elsewhere^14^. The first PCR cleanup was conducted with a Biometra^®^ Tpersonal ThermalCycler. Primer sequence details are given in the Supporting Information (SI Table S1). The PCR mixtures contained 1.5 μL of genomic DNA, 0.5 μL of amplicon PCR forward and reverse primers (10 μM), 2.5 μL of Taq Buffer 10x, 0.5 μL of Invitrogen dNTP (10 mM), 0.5 μL Titanium Taq 50x and 19 μL of sterile water. The final volume was 25 μL. The following cycling conditions were used for the amplification of DNA: initial denaturation at 95 °C for 3 min and 30 cycles of denaturation at 95 °C for 30 s, annealing at 55 °C for 30 s and extension 72 °C for 30 s, followed by a final extension at 72 °C for 5 min and a hold at 10 °C. PCR products were purified using 1.5 % agarose gel and with the GE Healthcare Kit (eluted with 20 μL of 10 mM Tris-Cl, pH 8.5). PCR purified products were quantified using the Quant-iT dsDNA HS assay kit and Qubit fluorometer (Invitrogen). The following steps were: second PCR clean up, library quantification and normalization, library denaturing and MiSeq samples loading. Illumina reads were processed using fastx_trimmer to remove barcodes (20 bp). PANDAseq with a quality threshold of 0.6 was used to assemble paired-end Illumina reads and raw sequencing data was processed and analyzed using the QIIME software package version 1.8.1. Quality filtered sequences were subsequently clustered into operational taxonomic units (OTUs) at a cutoff of 97% sequence identity using QIIME pick_closed_reference_otus.py script and the uclust algorithm. Taxonomic information for representative sequences for each OTU was performed with the Greengenes data base.

### 2.4 Statistical analysis

The microbial community structure was analyzed by comparing the similarity of the different microbial communities in the samples by principal component analysis (PCA). The analyses were performed in Excel XLSTAT statistical software and PCA used Spearman correlation matrix. The matrix was filled with the relative abundance of next generation sequence reads assigned to different microbial genera. Although 146 OTUs were identified, only 21 assigned OTUs were used to fill the matrix as they were the most representative groups of the 108 samples collected and this was to avoid overweighting the statistical analysis with insignificant OTUs.

The entire study area can be divided into samples that were never polluted nor treated (background = B), those that were polluted and treated so that the contaminant level was below detection limits (no longer polluted = N) and those that are still polluted (heavily polluted = SH; polluted with methane, which is a metabolite indicative of biodegradation processes = SM; polluted with both gasoline (benzene, toluene, ethylbenzene and xylenes – BTEX) and diesel compounds (polyaromatic hydrocarbons - PAHs) = S; polluted only by BTEX = SB, polluted only by PAHs = SP; polluted by BTEX and ethanol = SBE; polluted by ethanol = SE), as described in Table 2.

**Table 2.** Description of the state of groundwater samples from the field study.

| Samples | Description |
| --- | --- |
| B | Background: never has been contaminated or treated |
| N | No longer contaminated sites: BTEX, PAH and ethanol = below detection limit and methane < 5 mg L<sup>-1</sup> |
| SH | Heavily contaminated sites: BTEX > 5 mg L<sup>-1</sup> |
| SM | Pure biodiesel contaminated sites: methane ≥ 5mg L<sup>-1</sup> |
| SP | PAHs contaminated sites: above detection level |
| SB | BTEX contaminated sites: above detection level |
| SBE | BTEX and ethanol contaminated site: above detection level |
| SE | Ethanol-contaminated site: above detection level |
| S | PAH and BTEX-contaminated sites: above detection level |

## 3. RESULTS AND DISCUSSION

### Back to the wild

The different sources of pollution were cleaned-up using active and passive remediation strategies. As decontamination progressed, some microorganisms increased in relative abundance while others died out. These successional shifts are often conducive to the system recovery that trends towards the initial state^6,18^. Accordingly, the statistical evaluation of the perturbation caused by different parcels of pollution sources demonstrated that the local ecosystem was trending to the natural ecosystem equilibrium (Figure 2). The dynamics of microbial response to the perturbation was composed of three main clusters of samples from (1) the background, from (2) the no-longer-polluted and from (3) the still-contaminated sites. Heavily contaminated (SH) and BTEX-contaminated site (SB) samples clustered together since BTEX compounds were the main drivers for reducing microbial community resistance and altering their composition. Hydrocarbon degrading microorganisms were stimulated by the nutrient input provided by the contaminant and out-competed native communities unable to exploit the new resources. Although resilience was transiently hindered, these hydrocarbon degrading microbial specialists apparently degraded the pollutants and subsequently contributed to the overall system resilience by removing the pollution so that natural microbial communities were able to return to their initial levels. The no-longer-polluted samples clustered close to the background samples, which represent the initial ecosystem. Overall, microbial communities were able to reassert themselves and reestablish the ecosystem natural equilibrium.

**Figure 2.**
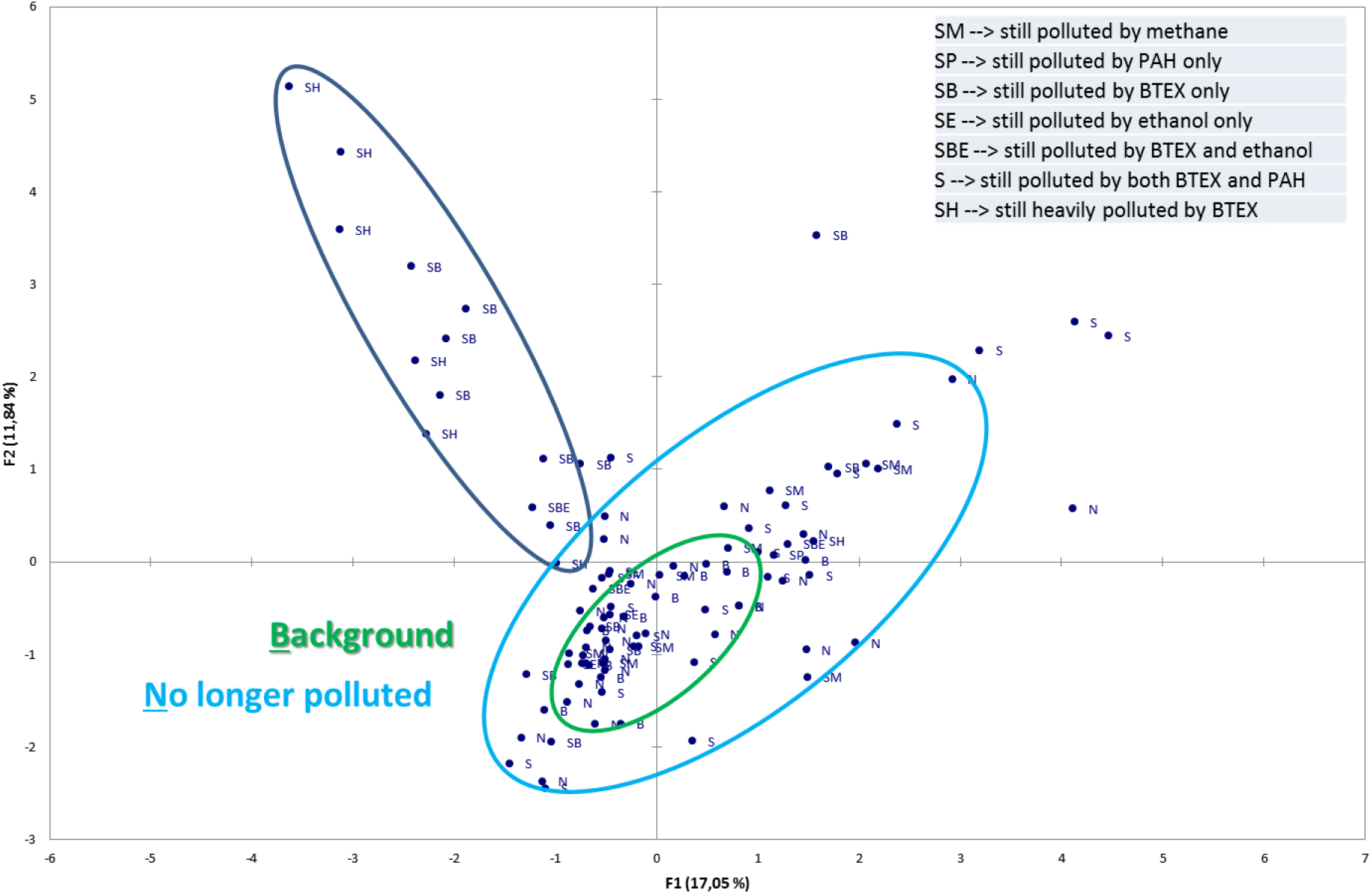
Principal component analysis (PCA) of the microbial communities in different samples taken over eight different pollution sources and remediation sites. The green ellipsoid encircles most of the background microbial communities and the light blue encircles most of the communities from groundwater that is no longer contaminated. The dark blue ellipsoid encircles communities from gasoline-contaminated groundwater whether heavily contaminated (SH), moderately contaminated (SB, SM, SP, S) or mixed with ethanol (SBE, SE).

### Identifying the players

Microbial community was composed by 146 identified genera, but only twenty-one genera presented predominant levels of sequence-assigned OTUs (operational taxonomic units) and thus, our discussion focused on these predominant genera. The evaluation of the relative abundance of certain taxonomic units (here at the genus level) as a function of the final state of the sample (B, N, or S) led to clear bioindicators for the pollution and subsequent ecosystem return towards the initial state (Figure 3). The shifts in microbial community profiles reflected the level and sources of pollution as well as the clean-up technologies applied. The microbial genera detected in heavily contaminated samples (SH) were dominated by *Telmatospirillum* spp., *Burkholderia* spp. and *Salinispora* spp. which are known hydrocarbon degraders^19,20 21 22–26^. Sites moderately contaminated were mainly composed of genera known to have specialist hydrocarbon degraders (e.g., *Rhodoplanes, Burkholderia, Gouta* 19, *Desulfurispora, Pelotomaculum, Desulfusporosinus* and *Geobacter*) and methanogens (*C. methanoregula* spp., *Methanosaeta* spp. and *Methanocella* spp.) as an indicative of the ongoing anaerobic biodegradation reactions. These pollution-selected microorganisms were the key contaminant-degrading agents and after the pollution pressure was alleviated, the surviving community was able to increase the original non-hydrocarbon degrading populations leading to the return towards the original (unpolluted) state. As the decontamination progressed, these pollution-selected microorganisms were replaced by microorganisms detected in background samples (*i.e*., *C. solibacter* and *C*. *koribacter*), demonstrating that the geochemical condition of the site could shape microbial composition.

**Figure 3.**
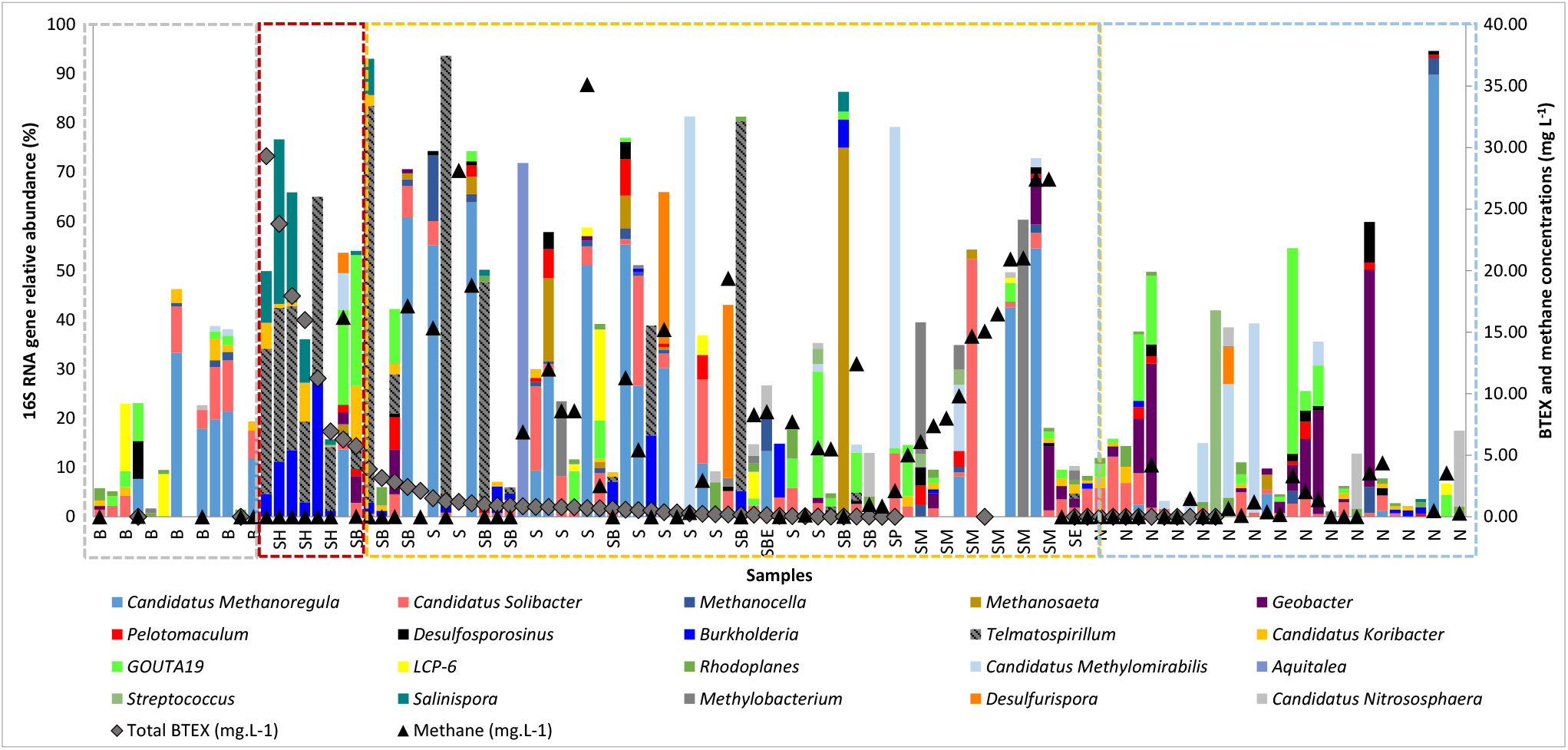
16S RNA average relative abundance (%) of indicator microbial groups and BTEX and methane concentrations (mg L^−1^) for different pollutant levels at multiple sites in the same groundwater.

### Passive vs active remediation technologies

Although the system converged towards its former state, the pollution sources and the clean-up technologies affected the rate of return to the pre-disturbed condition. The less intrusive technology was monitored natural attenuation (MNA) and it induced a faster return to the background condition relative to the active cleanup technologies (Table 3). For example, B20 releases under methanogenic (MB) and iron and sulfate biostimulation (ISB) have not yet returned to the microbial baseline condition, whereas for B20 under MNA, the perturbation lasted as long as the pollution and the return towards the natural equilibrium appeared to start shortly after the pollutants were degraded. A similar pattern was observed for B100 releases as MNA also increased (0.9 year) the return to the microbial background conditions relative to the partial chemical oxidation (PCO-B) treatment (1.7 years). Despite the initial community sensitivity to the pollution, MNA treatment induced minimum ecosystem disruption and led to the relatively rapid return to the initial microbial state.

**Table 3.** Overview of sites remediation state, length of treatment, microbial community return time to the unpolluted state (background) and total time for clean up treatment and biological recovery.

| Site | State* | Time for treatment (years) | Background return time (years) | Total time (years) |
| --- | --- | --- | --- | --- |
| B100 MNA | Decontaminated | 5.4 – 7.5 | 0.9 | 6.3 – 8.4 |
| B100 PCO-B | Decontaminated | 1.63 – 1.74 | 1.7 | 3.3 – 3.4 |
| B20 MNA | Decontaminated | 5.4 – 6.3 | 0 | 5.4 – 6.3 |
| B20 ISB | Decontaminated | 1.6 | NY | Greater than 1.6 |
| E85 MNA | Decontaminated | 3.2 – 5.3 | 0.72 | 3.9 – 6.0 |
| E25 NB | Still polluted | Greater than 10.9 | NY | Greater than 10.9 |
| E10 MNA | Still polluted | Greater than 5.8 | NY | Greater than 5.8 |
| B20 MB | Still polluted | Greater than 5.4 | NY | Greater than 5.4 |

*Notes.* NY: not yet returned to the background microbial profile.
\*Any of these sites have been entirely decontaminated, as there are still residual contaminants concentrations throughout the sites. This table refers exclusively to the samples included in this work.

### Pollutant dynamics in groundwater

The different dynamics undergone by biofuels in groundwater can substantially affect microbial community responses and the time required to decontaminate the site. Among the releases that were treated under MNA (E85, B100 and B20), the decontamination was faster (total time between 3.9 - 6 years) for the site polluted by gasohol release (E85) than for that contaminated by biodiesel blends (6.3 - 8.4 years for B100 and 5.4 - 6.3 years for B20). This can be attributed to the high solubility of ethanol that quickly dissolves into the groundwater and becomes more available to microbial attack as opposed to the viscous and less soluble biodiesel that tends to remain in the NAPL (non-aqueous phase liquids) (Ramos et al., 2016). Thus, biodiesel can last longer than ethanol and therefore, more intrusive cleanup technologies are generally applied to achieve remediation goals.

### Faster clean-up vs faster return to the baseline condition

Active clean-up technologies can influence microbial composition depending on both the level of pollution and the clean-up strategy applied. Partial chemical oxidation was applied to remove the poorly water-soluble biodiesel and prevent the long-term effects generally associated with source zones. The PCO-B treatment was successful in removing B100 contamination (after 2 years), but the return to the previous ecosystem condition was slower than the treatment of B100 by natural attenuation (MNA), since the PCO-B created rapid and relatively extreme effects on the local microbial communities. The same applied to the iron and sulfate biostimulation (ISB) that treated diesel-biodiesel release (1.6 years) faster than MNA (after 5 years), but for which the intensity of the disruption delayed the return to the background microbial profile. The longest effects resulted from gasohol blends with higher BTEX content (E10 MNA, E25 NB) and the B20 under methanogenic biostimulation (MB) in which acetate was added as supplementary substrate to produce methane in an anoxic environment. These sites are still polluted due to different factors that played a role (either pollutants and/or treatment) in driving the community even further from the natural state. The induced anoxic conditions certainly created a long-term ecosystem perturbation and impeded ecosystem recovery and site closure. These results demonstrated that the faster clean-up technology did not promote a faster return to background ecosystem conditions. This must be carefully accounted for in the remediation strategies’ decision-making process since rebuilding the natural ecosystem can be extremely long and expensive^2^. The information about each site microbial community profile is provided in the supporting information.

### Resilient responses

The site perturbation was ultimately followed by the return to the baseline microbial condition at most of the sites where the microbial communities were considered resilient. The identification of these resilient communities was conducted by the statistical evaluation of microbial communities in background and no-longer-polluted samples. The clusters were separated into samples that have returned to the initial microbial condition and samples that still contain signs of pollution bioindicators despite the removal of the chemical contamination (Figure 4). Samples that had not yet returned to the pre-disturbed biological condition clustered together with the pollution bioindicators. These bioindicators were the microbial genera *Geobacter, Pelotomaculum, Burkholderia, Desulfosporosinus* and some methanogens (i.e., *Methanosaeta, Methanocella* etc.). These genera where associated with relatively recent contamination^27–29^. Comparatively, the other cluster was composed mainly of background samples and some no-longer-polluted samples that likely returned to the pre-disturbed condition. These samples also clustered with a microbial assemblage that was mainly composed by *Koribacter, Methylomirabilis, Nitrososphaera* and *Solibacter*. Since these genera were detected at both background and no-longer-polluted samples and have not been closely associated with bioremediation reactions, their capacity to survive to the transient perturbation and to return to their predominant abundance shifted the microbial composition to the former state of the site before the perturbation.

**Figure 4.**
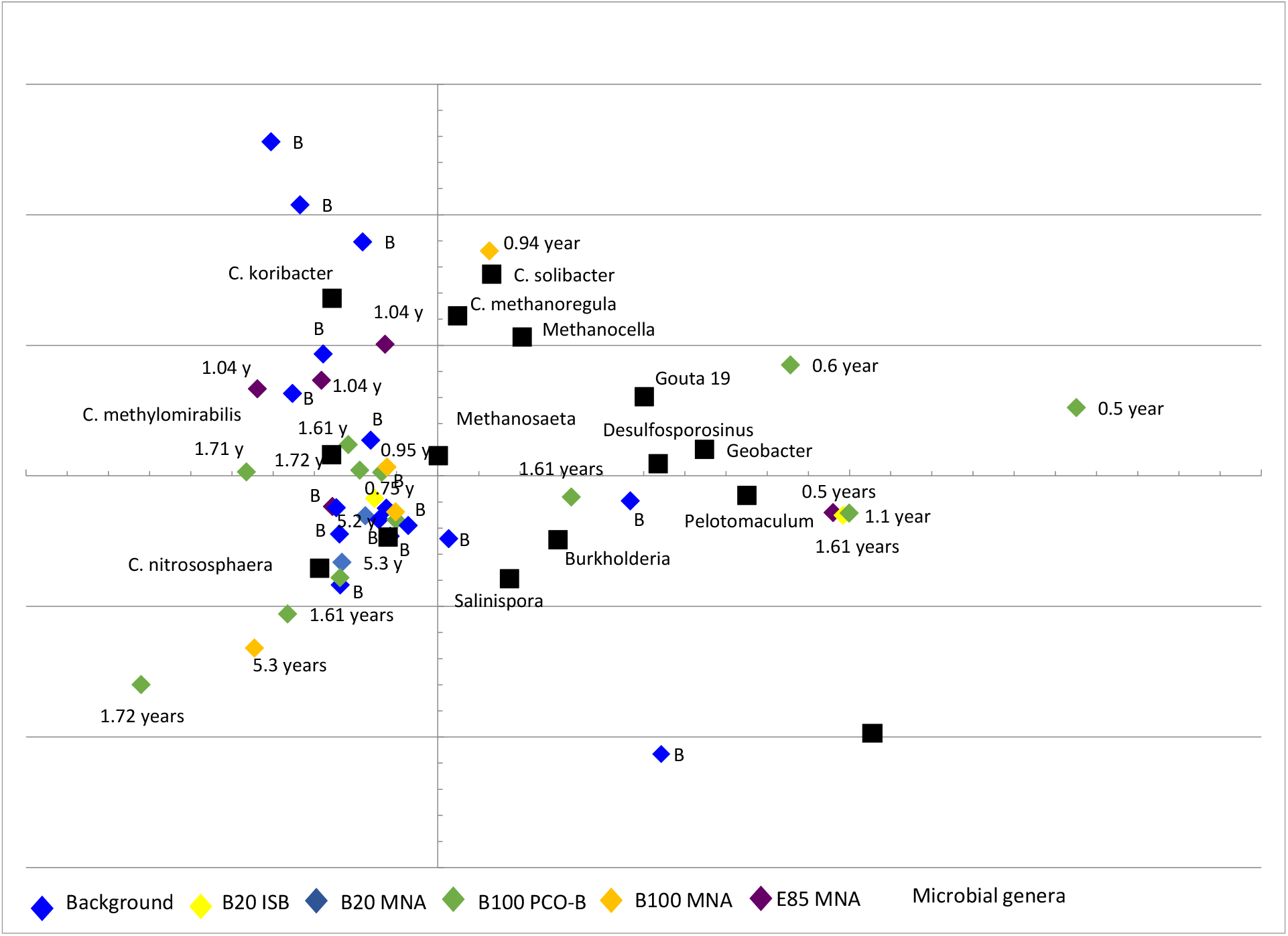
Principal component analysis (PCA) of the microbial communities in 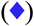 background and no-longer-polluted samples from B100 PCO-B 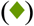, E85 MNA 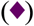, B100 MNA 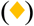, B20 ISB 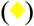 and B20 MNA 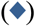 sites. Samples clustered in the left and right quadrants. Microbial genera (■) correspond to the main drivers for the final state of the samples. Right quadrant: no-longer-polluted samples that haven’t yet returned to the pre-disturbed microbial state. Left quadrant: background samples and no-longer-polluted samples that have returned to the pre-disturbed microbial state.

In summary, this long-term field study with multiple polluted parcels and treatment technologies over a homogenous groundwater aquifer demonstrated the relative resilience of the natural microbial community post pollution and treatment. Although the system converged towards its former state, the pollution sources and the clean-up technologies affected the rate of return to the pre-disturbed condition. The rate of return generally reflected the intensity of the clean-up treatments. Monitored natural attenuation promoted the faster return to the microbial background condition as opposed to the active clean-up technologies that speeded up the contaminants removal but delayed the return to the previous unpolluted microbial composition. These differences should be taken into account during the decision-making process on remediation strategies, depending on the ecosystem services desired to maintain or the risk posed to sensitive receptors.

## ASSOCIATED CONTENT

### Supplementary information

Microbial profile after pollution. Microbial profile at different sites (B100-MNA, B20-MNA, B20-MB, B100-PCO-B, B20-ISB, E10-MNA, E85-MNA and E25-NB). Schematic view of experimental area configuration (MNA control experiment). Primer sequences used for 16S rRNA sequencing analysis.

## AUTHOR INFORMATION

## AUTHOR CONTRIBUTIONS

D.T.R. conceived the conceptual development of the paper, run the bioinformatics analysis, interpreted the results, wrote the manuscript and co-wrote grant applications. H.X.C. conceived the conceptual idea of the paper, wrote grant applications and contributed to the discussions of the results. T.M.V. conceived the conceptual development of the paper, analyzed the data, revised the manuscript and led the overall project.

The authors declare no competing financial interest. All authors approve the publication.

### Funding Sources

This research was primarily funded by Petróleo Brasileiro S/A – PETROBRAS (contract number: 0050.0076426.12.9). Additional funds were provided by National Council for Scientific and Technological Development (CNPq) (special visiting researcher program – PVE contract number: 406061/2013-0) and the Coordination of Improvement of Higher Education Personnel (CAPES) (scholarship).

## ACKNOWLEDGEMENT

We thank Petróleo Brasileiro S/A – PETROBRAS, the National Council for Scientific and Technological Development (CNPq) and the Coordination of Improvement of Higher Education Personnel (CAPES) for providing the funding that supported this research. We thank Dr. Marilda Fernandes and Dr. Ana Maria Rubini Liedke for their help with the chemical and microbial analysis, respectively. We also thank Dr. Sandrine Demaneche for the Illumina Miseq sequencing. We dedicate this paper to the memory of Professor Henry X. Corseuil, colleague, mentor and friend.

